# The IGVF Data Portal: a repository for insights into genome function

**DOI:** 10.64898/2026.11.18.767169

**Authors:** Jennifer Jou, Zhewei Shen, Khine Zin Lin, Shengcheng Dong, Ian Whaling, Otto Aleksi Jolanki, Pedro Assis, Idan Gabdank, Keenan Graham, Meenakshi Kagda, Mingjie Li, Forrest Tanaka, Lars Steinmetz, Ben Hitz

## Abstract

The Impact of Genomic Variation on Function (IGVF) Consortium has utilized a wide variety of assays and computational methods to study how genomic variation impacts function. To support the collaborative efforts of labs within IGVF, and to share the results of these efforts, the IGVF Data Administration and Coordination Center (DACC) has developed a data portal (https://data.igvf.org/) that serves as the repository for all results from the IGVF Consortium and as the source of results that have been integrated into the IGVF knowledge graph. Data generated by the IGVF Consortium, which has reached over 240 terabytes as of November 2026, can be publicly accessed via web browser, or programmatically via API, as well as via agentic tools. Here, we provide a summary of the available IGVF data and an overview of the design of the IGVF Data Portal.

## Introduction

The Impact of Genomic Variation on Function (IGVF) Consortium began in 2021 to investigate how genomic variation in both coding and non-coding regions affects function, building off of numerous other earlier and ongoing efforts to study genome function. Consortium members study this same question using mapping, perturbation, and predictive methods, and integrate these different approaches with one another to enhance our understanding of how genomic variants affect cellular phenotypes and, eventually, organismal phenotypes **[1]**.

Within IGVF, one of the Data Administration and Coordination Center (DACC)’s key goals was to develop the IGVF data resource, a database and data portal that stores and curates IGVF results before making them publicly accessible to the scientific community. As of November 2026, this database now includes over 100,000 publicly accessible files corresponding to over 240 terabytes of data from 65 assays, in both mouse and human samples, and 18 computational prediction methods, with more data anticipated through May 2027.

The data portal and its underlying model are designed to accommodate the highly varied IGVF data, which spans experimental assays, computational predictions, and more, allowing all of the data to be brought together into one centralized repository, which can be accessed both programmatically or via a user friendly, human-navigable web portal at https://data.igvf.org/. The portal also serves as the repository for the data that is loaded in the IGVF Catalog. All data in the portal, whether they are experimental or computational results, are accompanied by rich metadata about the experiment or method that adheres to defined, structured formats in accordance with FAIR (findable, accessible, interoperable, reusable) principles for data management **[2]**. In doing so, the DACC ensures that the Consortium can efficiently disseminate IGVF products and increase the impact and utility of IGVF data.

## Results

### Data on the Portal

As of November 2026, the IGVF Data Portal features 1,113 experimental analyses (deriving from 7,938 raw data sets) and 433 computational predictions. In total, there are over 100,000 raw and processed files, corresponding to over 240 terabytes of data. These data were generated by 113 IGVF labs across 26 awards and 7 affiliate labs.

The portal features experimental data from 65 unique assays in total, which can be divided broadly into mapping and characterization categories. The mapping category includes 36 types of assays, 24 of which utilize single-cell technology. These assays cover several different modalities, including chromatin accessibility, transcriptomics, imaging, methylation, chromatin structure, protein interaction, DNA binding, and spatial transcriptomics. The characterization category includes 14 types of CRISPR screen, 8 types of reporter assays, and 7 protein scanning assays. The assay categories are designed to highlight the key features of an IGVF study and are not mutually exclusive, so a single experiment on the portal often belongs to several categories at once. For example, a 10x Multiome experiment is grouped under single cell, chromatin accessibility, and transcriptomics.

The assays detailed above cover a wide breadth of biological space. The portal currently tracks over 12,000 biosamples, divided into over 8,900 human samples, 3,200 mouse samples, and 38 mixed human-mouse samples. These samples can also be divided according to classification: over 4,600 tissues or organ samples, 4,600 cell line samples, 2,100 primary cells, 1,200 differentiated cell specimens, 120 cell cultures, 90 organoids, 50 reprogrammed cell specimens, and 5 gastruloids (**Figure 1**). There are a further 76 yeast cell samples used in yeast two-hybrid assays. Many of these samples are additionally multiplexed using either barcode-based or genetic multiplexing methods; others are pooled in cell villages. Attribution metadata is tracked for samples so that sample sharing between IGVF labs is recorded. In addition to the experimental results, the portal also features predictive results from 18 computational methods. These predictions can be categorized into 11 major categories, shown in **Table 1**.

**Figure 1:**
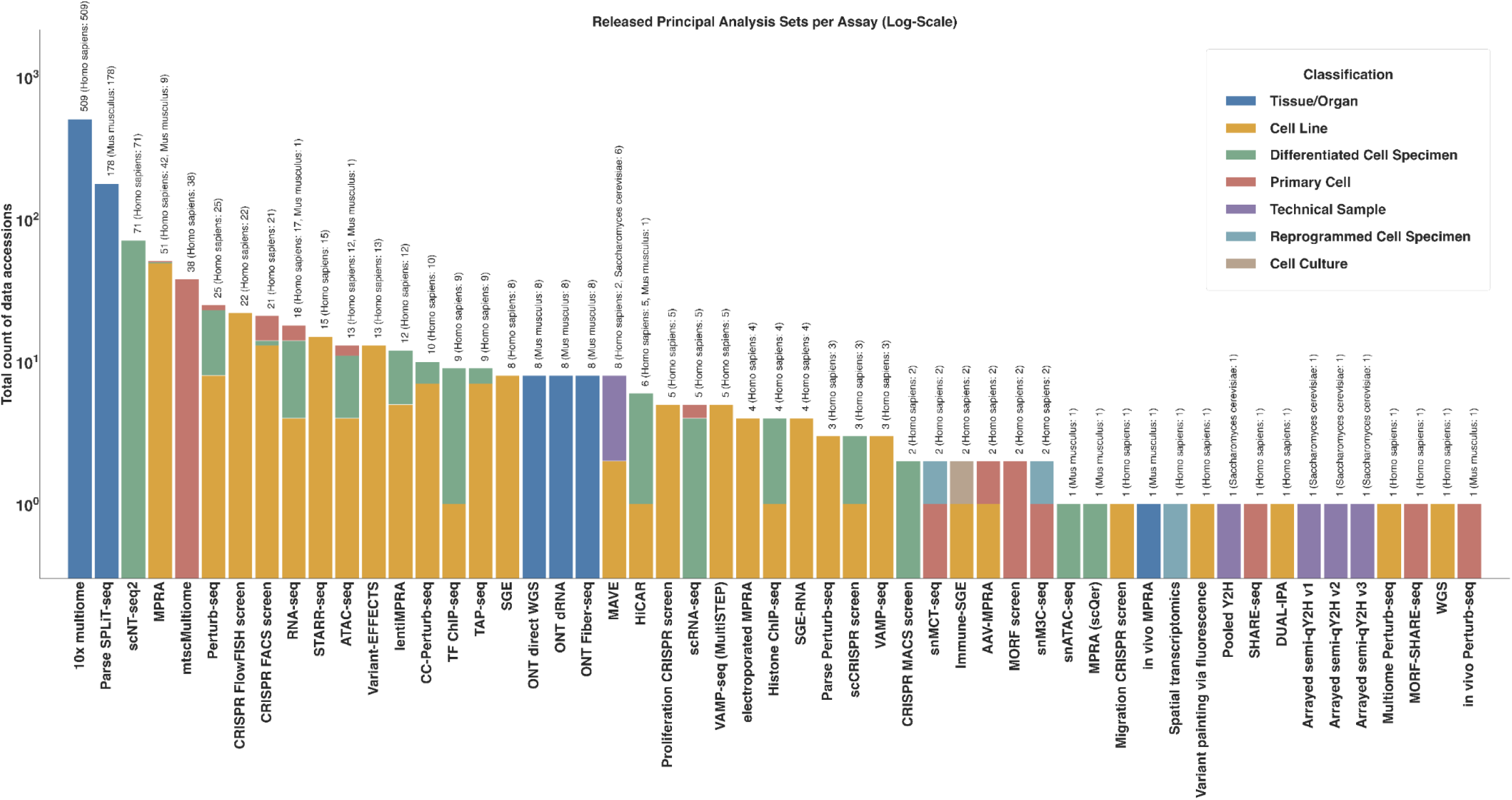
Summary of released experimental data and analysis on IGVF Data Portal. A stacked bar chart in log 10 scale, indicating the number of released principal Analysis Sets available on the IGVF Data Portal as of November 2026 per assay title. Each bar is subdivided by the classification of sample assayed, and is additionally annotated with details on the taxa of the samples. The data represented in this figure can also be viewed at https://data.igvf.org/search/?type=AnalysisSet&file_set_type=principal+analysis&status=released.

**Table 1:**
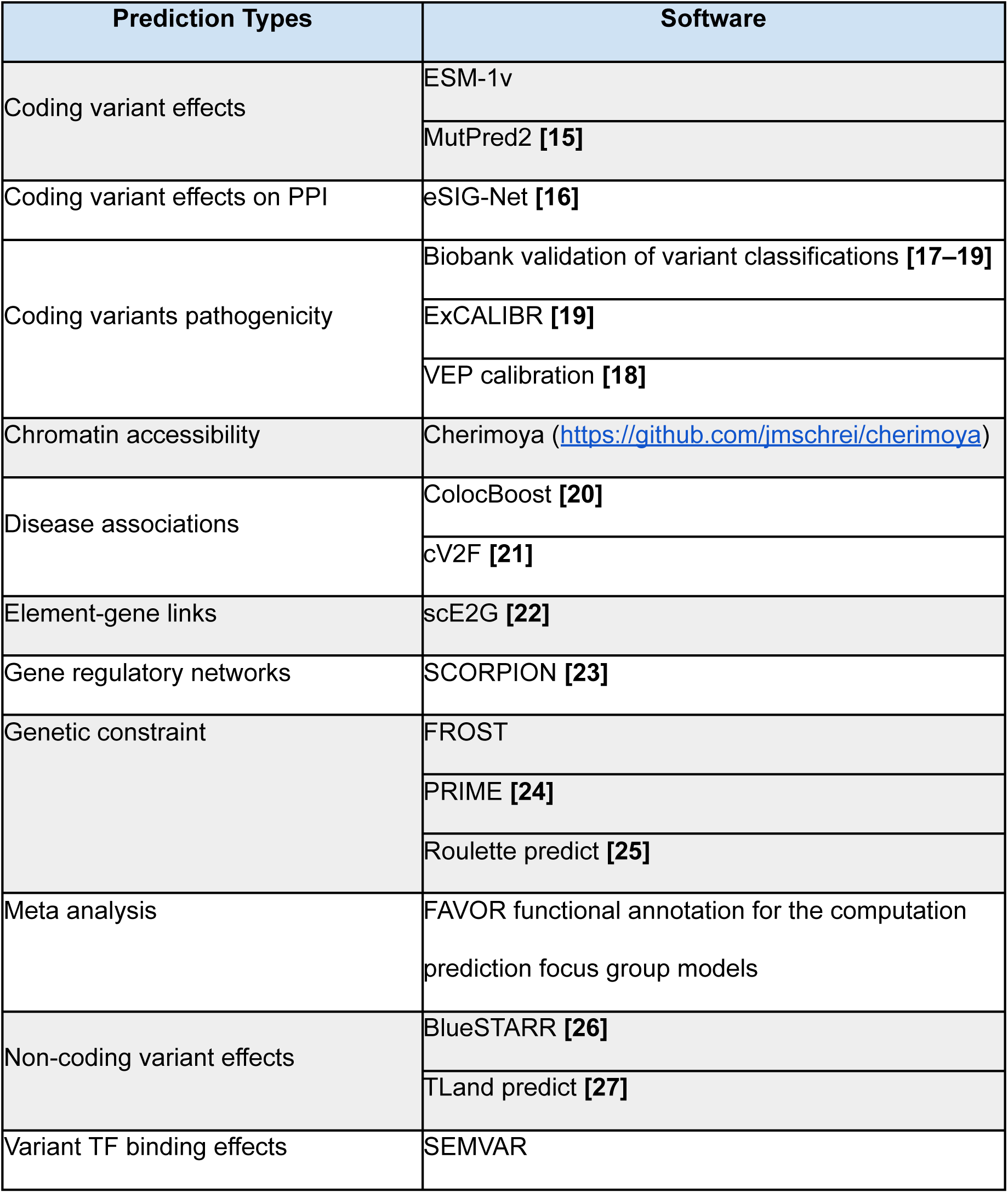
Prediction data on IGVF portal. Table 1 lists all the prediction software and the prediction output data content that have been released on the IGVF data portal as of November 2026.

| **Prediction Types** | **Software** |
|---|---|
| Coding variant effects | ESM-1v |
|  | MutPred2 **[15]** |
| Coding variant effects on PPI | eSIG-Net **[16]** |
| Coding variants pathogenicity | Biobank validation of variant classifications **[17–19]** |
|  | ExCALIBR **[19]** |
|  | VEP calibration **[18]** |
| Chromatin accessibility | Cherimoya (https://github.com/jmschrei/cherimoya) |
| Disease associations | ColocBoost **[20]** |
|  | cV2F **[21]** |
| Element-gene links | scE2G **[22]** |
| Gene regulatory networks | SCORPION **[23]** |
| Genetic constraint | FROST |
|  | PRIME **[24]** |
|  | Roulette predict **[25]** |
| Meta analysis | FAVOR functional annotation for the computation prediction focus group models |
| Non-coding variant effects | BlueSTARR **[26]** |
|  | TLand predict **[27]** |
| Variant TF binding effects | SEMVAR |

In addition to new results from IGVF labs, the IGVF Data Portal stores reference files and supplemental files that provide context to those results. For example, copies of the designated IGVF standard genome and transcriptome reference files from GENCODE, with metadata to track the source of these external files, are available via the data portal. Labs also provide supplemental lists of variants and guide RNAs or oligos that are used to analyze experimental data.

The IGVF Consortium releases data on a rolling basis as data is finalized, rather than on a fixed schedule, with the goal of making data available to the public as soon as possible. All submitted data go through a combination of automated checks and human review prior to release to ensure their quality.

### Design of the Portal Database

The IGVF database is based upon SnoVault, a general-purpose, hybrid object-relational database framework used to manage, store, and serve large-scale genomic metadata **[3]**. SnoVault enables flexible schemas and schema validation using JSON-LD (https://json-ld.org/) and JSONSchema (https://json-schema.org/). The database is deployed on AWS RDS (Postgres, relational database), and AWS OpenSearch is used to operate an object database for search and caching. The frontend of the data portal is a nextJS application deployed to AWS Fargate, while the backend is a Python application also deployed to AWS Fargate. Both the frontend and backend services are set up to automatically scale to respond to demand, ensuring a smooth user experience and efficient resource utilization (**Figure 2**). The code for the IGVF portal backend (https://github.com/IGVF-DACC/igvfd) and frontend (https://github.com/IGVF-DACC/igvf-ui) is open source and available on GitHub.

**Figure 2:**
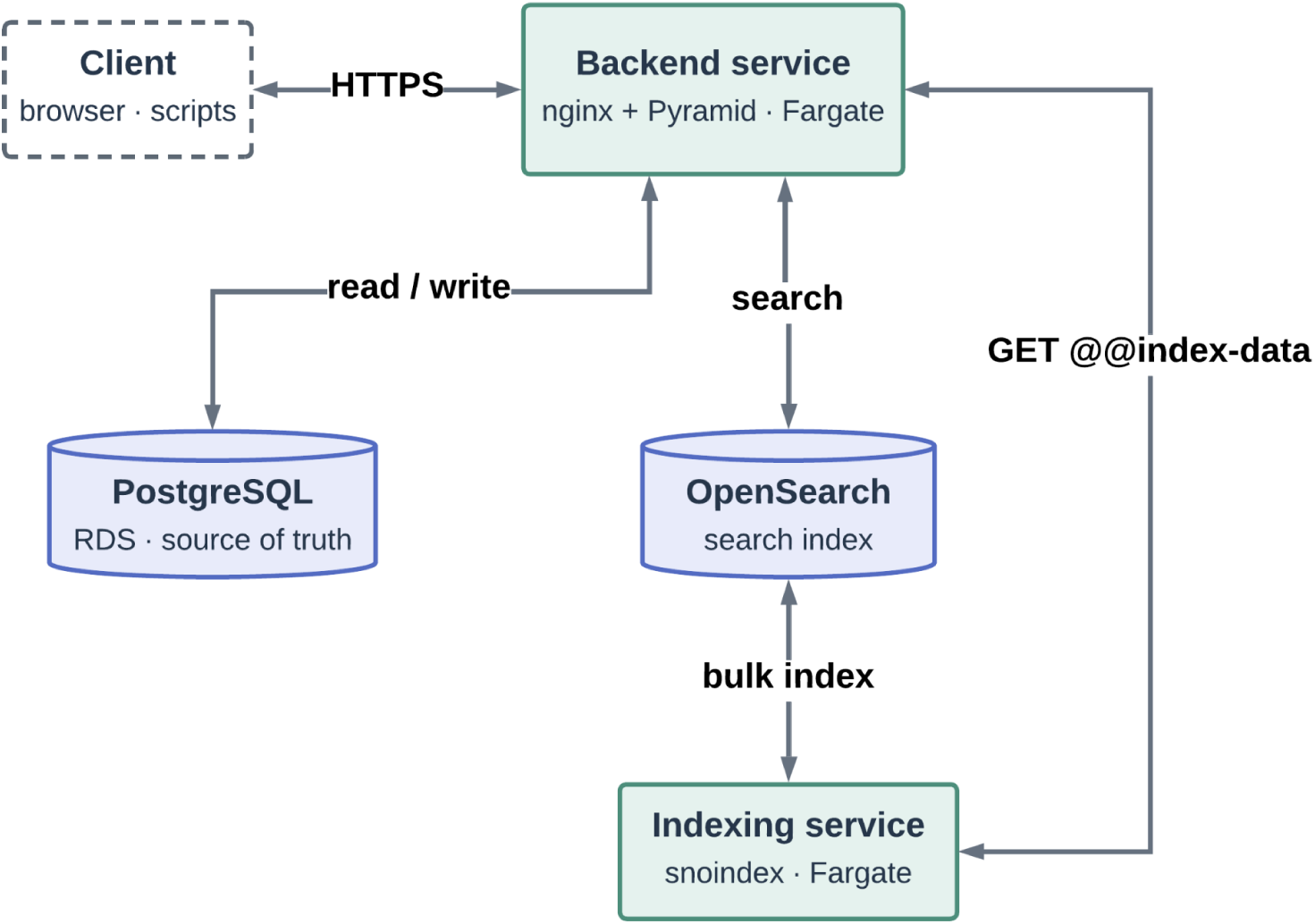
Overview of SnoVault. This diagram shows the request flow through the IGVF Data Portal. Browsers load pages from igvf-ui (Front end), whose Next.js server renders React and fetches JSON-LD from igvfd (Backend). The browser then calls igvfd directly for client-side updates. Scripts and submission tools use the same igvfd REST API without going through the UI. The igvfd application writes to PostgreSQL, the source of truth, and serves search and embedded documents from OpenSearch, which the indexing service keeps current.

Data file storage is primarily via AWS S3. The IGVF Consortium also does periodic data transfers of released data to NHGRI Analysis Visualization and Informatics Lab-space (AnVIL) as an additional data repository, and URLs to the corresponding AnVIL file location are stored in each file’s metadata **[4]**. The genomic data sharing consent status is tracked for each file on both the IGVF Data Portal and AnVIL sides to ensure access is blocked for controlled access data. Released IGVF data, including controlled access data, are also available for the scientific community on dbGaP (https://dbgap.ncbi.nlm.nih.gov/beta/study/phs003472.v1.p1/#study) with periodic updates. In addition, DOIs are minted daily for newly released IGVF datasets using a DACC-Crossref (https://www.crossref.org/) submission service, providing persistent, citable references to the data.

### Data Modeling and Organization

At a high level, the data model, which relies on the previously mentioned JSON-LD framework, divides data into broad categories of experimental raw data, processed data, pseudobulk data, and prediction data. The schema allows for linkages between all types of data in order to reflect how IGVF labs use IGVF data to generate new predictions or pseudobulks of single cell data, which can also be used as prediction inputs. In doing so, the DACC emphasizes the discoveries made possible via collaboration across IGVF labs.

One IGVF Consortium collaborative effort that connects these different types of data is the element to gene (E2G) prediction project. In this project, mapping centers within the Consortium first generated single cell transcriptomic and chromatin accessibility data. These were then processed using a uniform single cell processing pipeline developed by IGVF labs (manuscript in preparation). The single cell processing results were annotated by Consortium members, and based on those annotations, pseudobulked data sets were generated. Lastly, these pseudobulked data were used as input for single cell E2G methods, shared on the portal as prediction sets. Data from each step of this process, from start to finish, are stored on the portal with metadata showing how these inputs and outputs relate to each other, helping users understand the data provenance and also view detailed information about each analysis done. In a similar case, the ChromBPNet **[5]** prediction effort also integrates single cell chromatin accessibility data with pseudobulking to produce models and predictions on how sequence variants affect chromatin accessibility.

At a more detailed level, the data model is composed of 56 object types, also referred to as “schemas,” that represent various components of an experiment. These objects are listed comprehensively on the portal’s schema page (https://data.igvf.org/profiles/), where users can view the many metadata properties defined for each object. The strictly defined schemas, with limited free text properties, ensures that metadata is captured in a regularized manner, which is more readily machine- and AI-readable. Some of these properties track static metadata, such as the file format of a file; other properties are defined as links to other objects within the database. This allows a file to be associated with a file set, a sample to be associated with an ontology term, or a donor to be associated with a sample. Object embeddings and calculated properties, which are populated automatically by the portal application, allow a limited degree of denormalization where appropriate, so that users viewing a sample will see the linked ontology related metadata without viewing the ontology term object directly.

The key concepts within the data model are files, file sets, and samples. Files represent individual data files, file sets organize related files and metadata associated with experimental or computational activities, and samples represent the biological material used in experiments or the biological context applicable to a computational prediction. These 3 object types, in addition to donors and workflows, are accessioned objects, meaning they receive an immutable and permanent accession identifier. Supplementing the key objects are other schemas used to capture metadata that contextualizes each experiment, such as treatments, modifications, software, and more. Importantly, ontology terms from 13 ontologies, listed at https://data.igvf.org/help/general-help/ontologies/, are included in the data model in order to annotate samples, assays, platforms, and phenotypes in order to maximize interoperability with other data resources.

The file, file set, and sample objects are all subclassed in order to better handle the diverse nature of IGVF data. For example, file sets are divided into 8 subclasses representing different components of an assay or computational analysis: measurement sets, auxiliary sets, analysis sets, construct library sets, curated sets, model sets, prediction sets, and pseudobulk sets, with additional metadata about the file set type for further subcategorization. File sets can be linked modularly in a flexible, many-to-many fashion, which accommodates cases in which raw data in a measurement set is processed more than once, producing different sets of processed data that are each grouped into an analysis set. Analysis sets may also link to each other to represent multi-step analyses, such as aggregation across replicates for the purpose of cell type annotation. Similarly, the construct library set, which captures details about guides or oligos used in CRISPR-based assays and reporter assays, can be associated with multiple samples, representing instances where the same set of constructs were used in more than one experiment. Meanwhile, the sample object is also split into 6 subclasses. One of these is the multiplexed sample, a schema designed to capture sample multiplexing metadata, which is needed to handle the widespread adoption of these techniques in genomics to enable more cost efficient and higher throughput sequencing.

**Figure 3** illustrates example data models used to represent three types of results, among many others, on the data portal. In **Figure 3A**, a mapping experiment using the Parse SPLiT-seq assay is divided across multiple file sets: the measurement set for the raw data, the intermediate analysis set for the multiplexed results, and the principal analysis set downstream of the intermediary results that have been merged, demultiplexed, and then regrouped on a per donor or sample basis. **Figure 3B** is an example of a Perturb-seq characterization experiment. Here, a construct library set stores metadata associated with the guide RNAs and an auxiliary set stores sequencing data on the gRNAs, both of which are associated with but independent of the measurement set, which has the sequencing data for the readout of the Perturb-seq assay. **Figure 3C** features an example of a computational prediction. Trained models are kept separate in a model set, with prediction results belonging to a prediction set, analogous to the relationship between a measurement set and an analysis set. Links to the software and genes provide additional information on the methodology used and the scope of the prediction.

**Figure 3:**
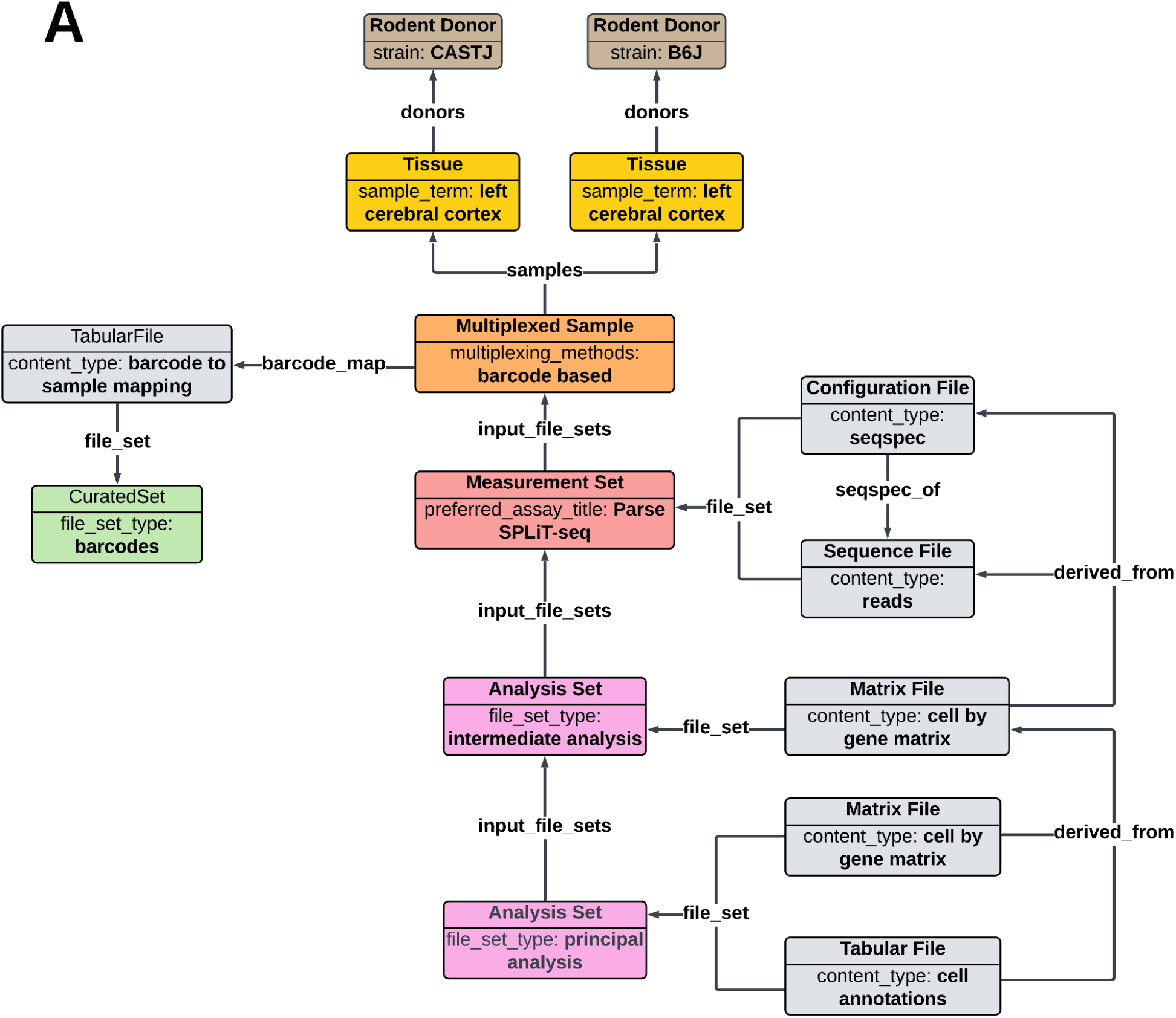
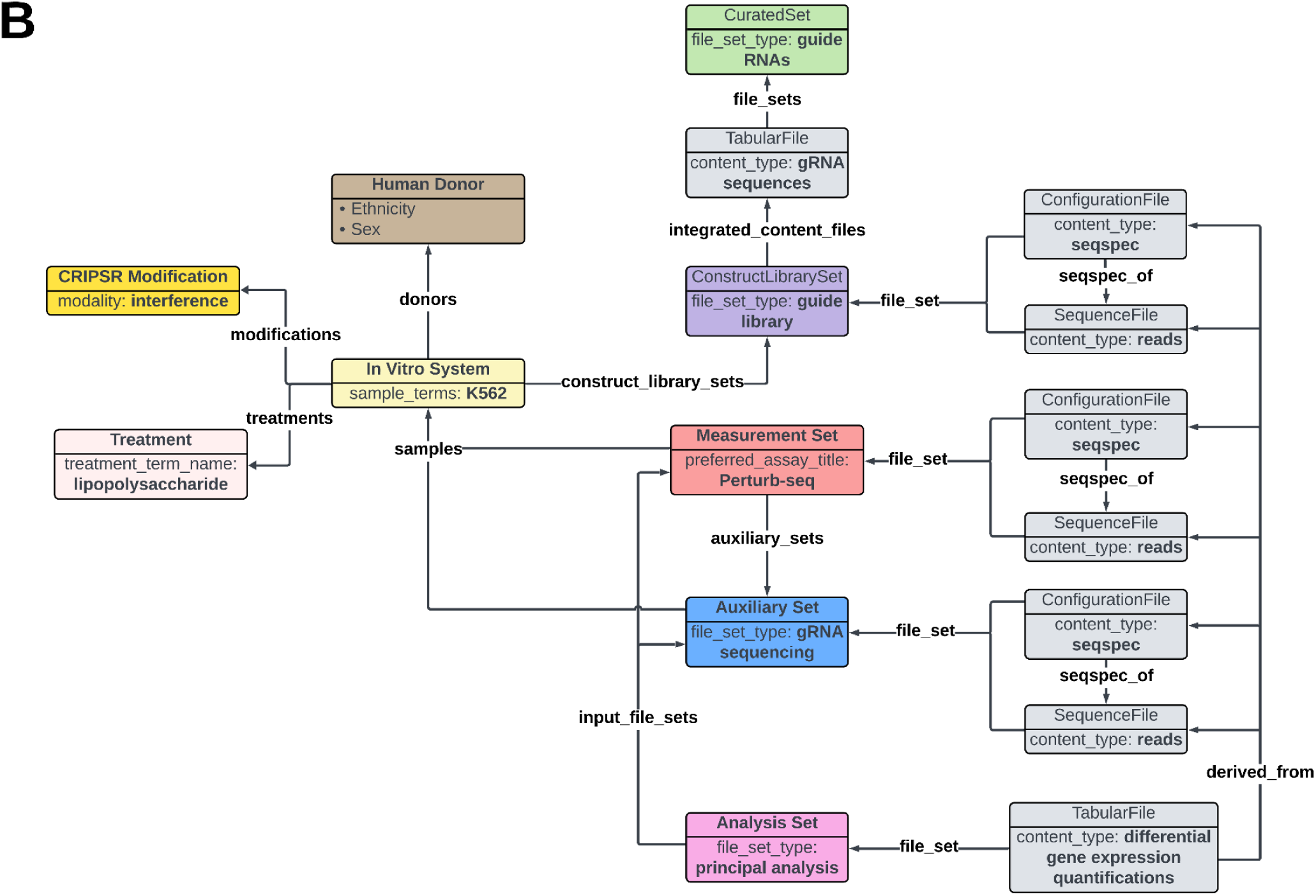
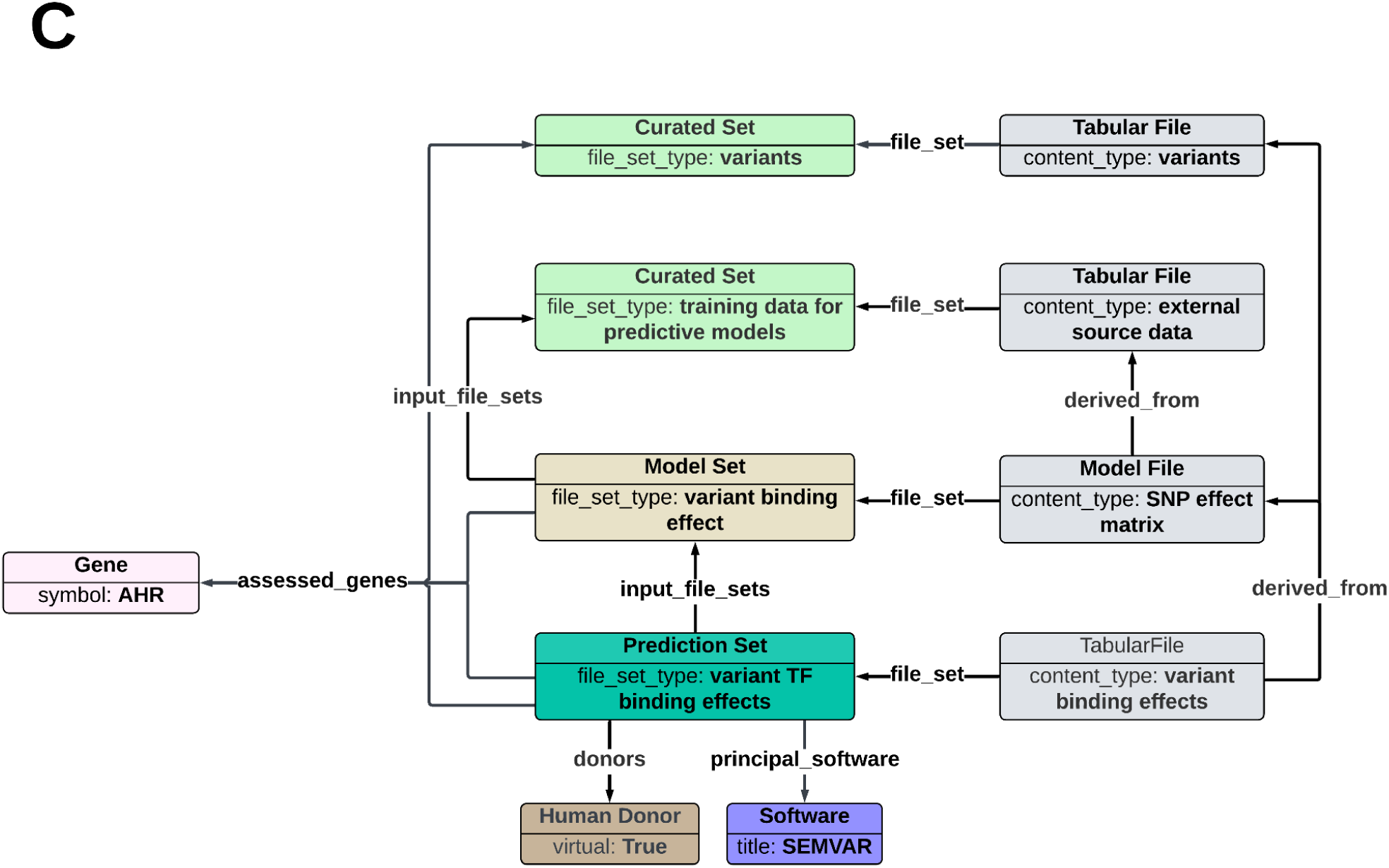
Examples of the IGVF Data Portal object types and data models. **(A)** A diagram showing an example data model for a Parse SPLiT-seq mapping experiment. The multiplexed sample, a subclass of samples, is used to group together the input samples in the multiplex. It can also link directly to a tabular file providing the mapping of barcodes to samples. The measurement set contains raw data files, such as FASTQs and seqspec files. The first intermediary stage of analysis is stored in a downstream intermediate analysis set, while the final stage of analysis containing demultiplexed data is stored in the furthest downstream principal analysis set. **(B)** A diagram showing an example data model for a Perturb-seq characterization experiment. Guide RNA metadata is represented by the construct library set, which is associated with the sample in which the constructs were transfected. The auxiliary set stores data from the gRNA sequencing so that it is separate from the sequencing data associated with the Perturb-seq readout. A principal analysis set is downstream of both the measurement sets and auxiliary sets. **(C)** A diagram showing an example data model for a computational prediction using the SEMVAR method. The model set contains the model file while the downstream prediction set contains the output of the model – in this case, scores indicating the effect of the variant on the binding affinity of transcription factors. Both the model set and prediction set indicate the assessed genes so that users can search for predictions according to a gene of interest. The prediction set also links directly to the relevant software method used to produce the prediction.

The IGVF data model is a “living” data model that the DACC updates on a weekly basis to accommodate new types of data from consortium labs. Weekly changes are reported on the Release Notes page https://data.igvf.org/releases/. This system increases the flexibility of the model and allows the database to quickly adapt to changing needs.

### Metadata and Data Validation

Beyond merely providing access to data, the DACC also endeavors to ensure the data presented to the scientific community are reusable and of high quality. To do so, a system of audits and file validation tools are used to review the quality of the metadata and data, respectively.

To review metadata quality, the IGVF application includes an auditing system within it that automatically inspects all objects in the database according to DACC defined guidelines. These audits are categorized by severity, with the two most severe categories being considered release blocking and requiring remediation by the submitting labs; the less severe categories may still be released depending on lab and DACC discretion. The audits can inspect linkages between objects and identify unexpected or missing metadata properties; for example, audits are used to check if cell line samples are linked to the correct donor object. Audits are also used to flag files that are missing a file format specification document. Any errors found via audit are automatically viewable on the data portal so that users are made aware of any potential issues with the data. A comprehensive list of implemented audits and their descriptions is available at https://data.igvf.org/audits/.

For data quality, the DACC maintains a file validation tool, referred to as “Checkfiles” (https://github.com/IGVF-DACC/checkfiles), to run automated checks. The tool relies on several validation tools developed by other groups. These data validity checks, which occur when a file is uploaded to the data portal, vary in extent depending on the type of the file. All files are checked for file corruption errors. Other files, such as FASTQs, BAMs, and BED, among others, have content checks to ensure there are no errors such as quality string length discrepancies, coordinates or contigs not present in the Consortium’s standard genome reference, and so on.

Certain tabular files, as categorized by their content types, have further specific content expectations with Consortium-defined formats developed by different IGVF focus groups, which are described at https://data.igvf.org/methodology/computational_standards/. Consortium members are expected to adhere to these formats. To systematically review these files for correctness, the DACC took these formats and implemented them within Checkfiles, allowing the tool to identify issues with incorrect value types or headers. For example, the CRISPR focus group developed the guide RNA sequences file format, while the MPRA focus group developed standardized formats for oligo sequence design tables and several types of MPRA analysis files. These efforts have improved the uniformity and interoperability of IGVF results and simplified data ingestion and interpretation into the IGVF Catalog Knowledge Graph database.

### Data Navigation

Navigation on the IGVF portal begins at the portal’s front page, which displays high level statistics about the total released data available at the portal. A sidebar provides links to the rest of the site, divided into five categories (**Figure 4A**). The Data section contains links to file sets and files. The Resources section contains various informational pages: assay by sample data matrices, informational pages on the IGVF grants, tables of IGVF software products, experimental and computational standards, lists of reference files used in IGVF, and details on the database auditing system. The Data Model section includes links to the JSON schemas and examples of how different types of experiments are modeled. The About section includes general information about the IGVF Consortium and its policies. Lastly, the Help section links to various documentation and help pages for both IGVF submitters and general users, as well as a link to the IGVF help desk email for questions and suggestions. Future portal updates will include front page changes to feature key resources and IGVF products.

**Figure 4:**
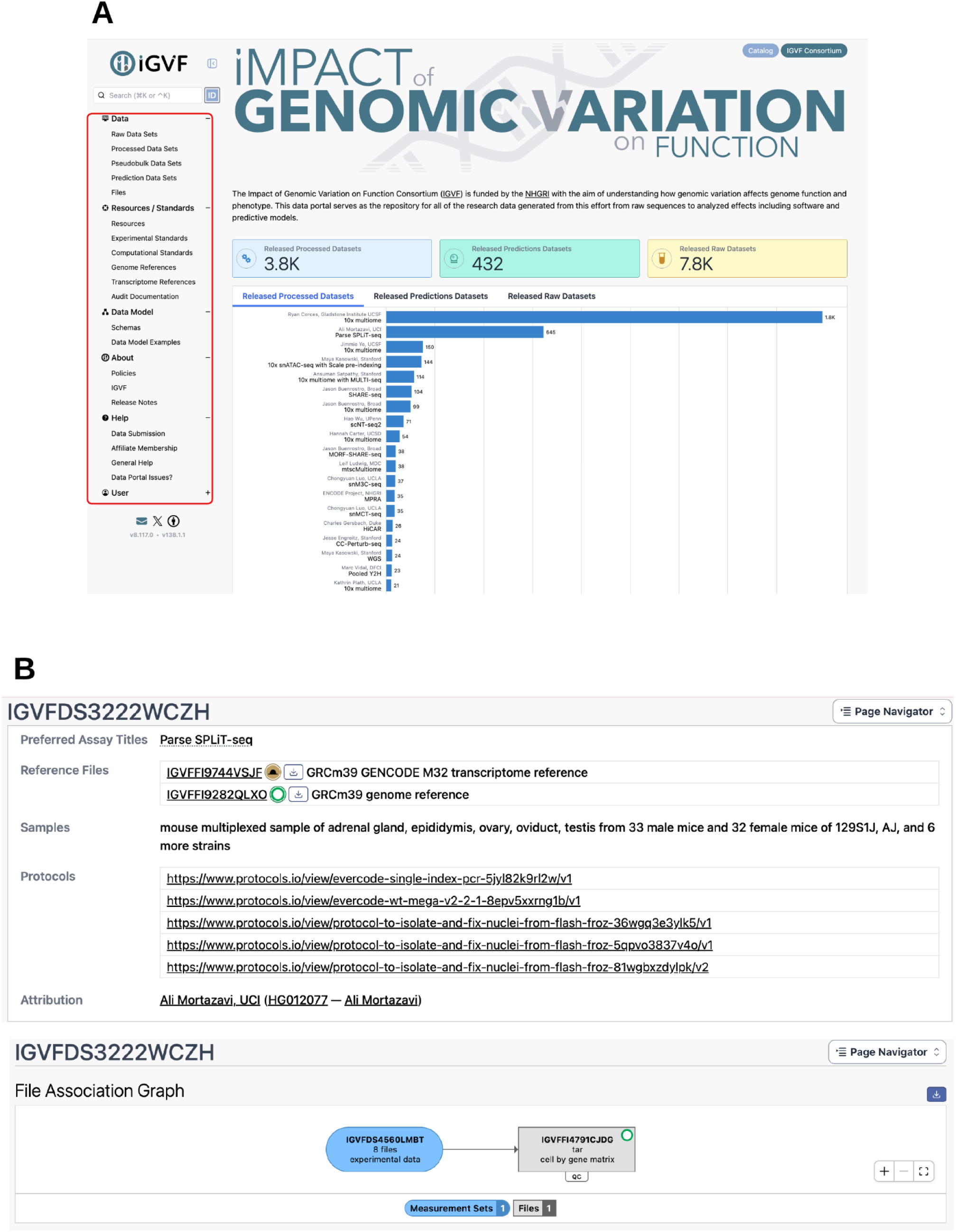
IGVF Data Portal navigation. **(A)** Portal home page and navigation. A screenshot of the IGVF portal’s home page, which features charts highlighting the number of released file sets. The sidebar, circled on the left, provides access to the rest of the portal, including file set search, data model information, documentation, and more. **(B)** Data object display example. A screenshot of an analysis set page on the IGVF portal. Basic summary information is displayed at the top of the page, such as metadata regarding the assay and a sample summary. Below are the file association graph and tables of the files, samples, donors, and related file sets.

All objects in the database have an individual summary page that displays metadata about the object. For example, for a typical analysis set, the summary page displays a table of files generated by that analysis, a file association graph visually illustrating the provenance of files, tables of associated samples, and other useful information on the sample (**Figure 4B**). The page also includes links to upstream file sets so that users can trace back and view the provenance of the processed results.

To help users locate data of interest, the data portal includes two search views: a default list view and a report view that is suited for viewing and downloading tables of metadata (**Figure 5**). Either view can be filtered using the “facets” located in the sidebar, allowing users to quickly refine the search results according to assay, sample, lab, and many other metadata properties (**Figure 6**). By default, a selection of facets are automatically displayed on all search result pages, chosen by the DACC to highlight metadata fields that are commonly used. Users can also customize their search experience by choosing additional optional facets for other metadata fields, tailoring the sidebar based on their specific needs. Facets can also be reordered so that users can place their most used facets at the top of the page; the custom layout is then saved to the user’s account or in the web browser cookies and retained upon future visits to the portal. This system allows the portal interface to remain uncluttered while also providing flexibility for more targeted and refined searches.

**Figure 5:**
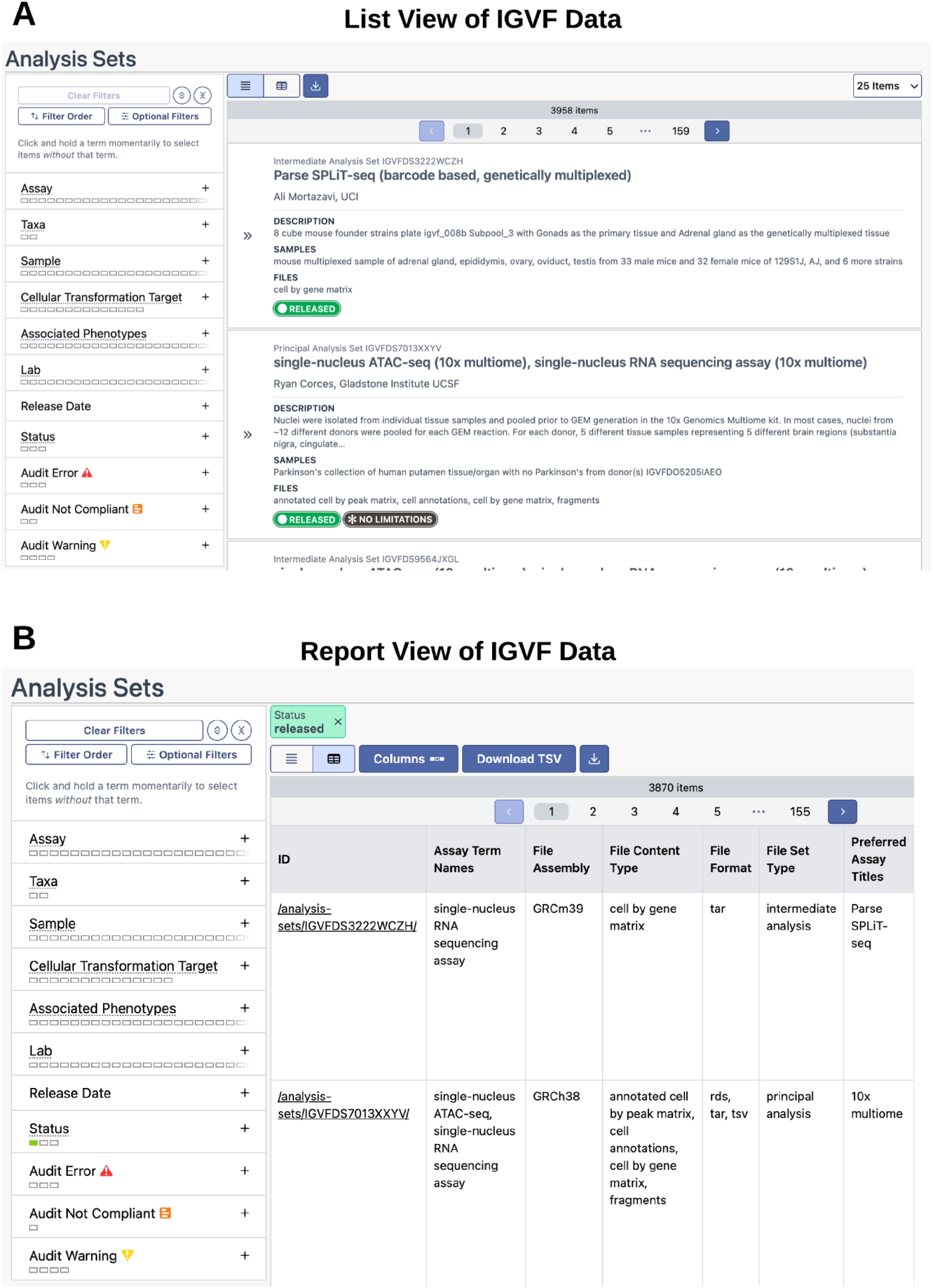
IGVF data query. There are two methods to display query results on the IGVF Data Portal, with results being displayed as either item cards or a table with selectable metadata columns. (**A)** Search list view. The search list view style returns query results as a list of data objects with the key metadata such as description, samples, and types of files displayed in each item card. (**B)** Report list view. The report list view returns query results as a table with metadata columns. Some of the most frequently used metadata are displayed as the default columns when switching to this view style. The “Columns” function allows users to include or exclude all available metadata associated with query results in this table.

**Figure 6:**
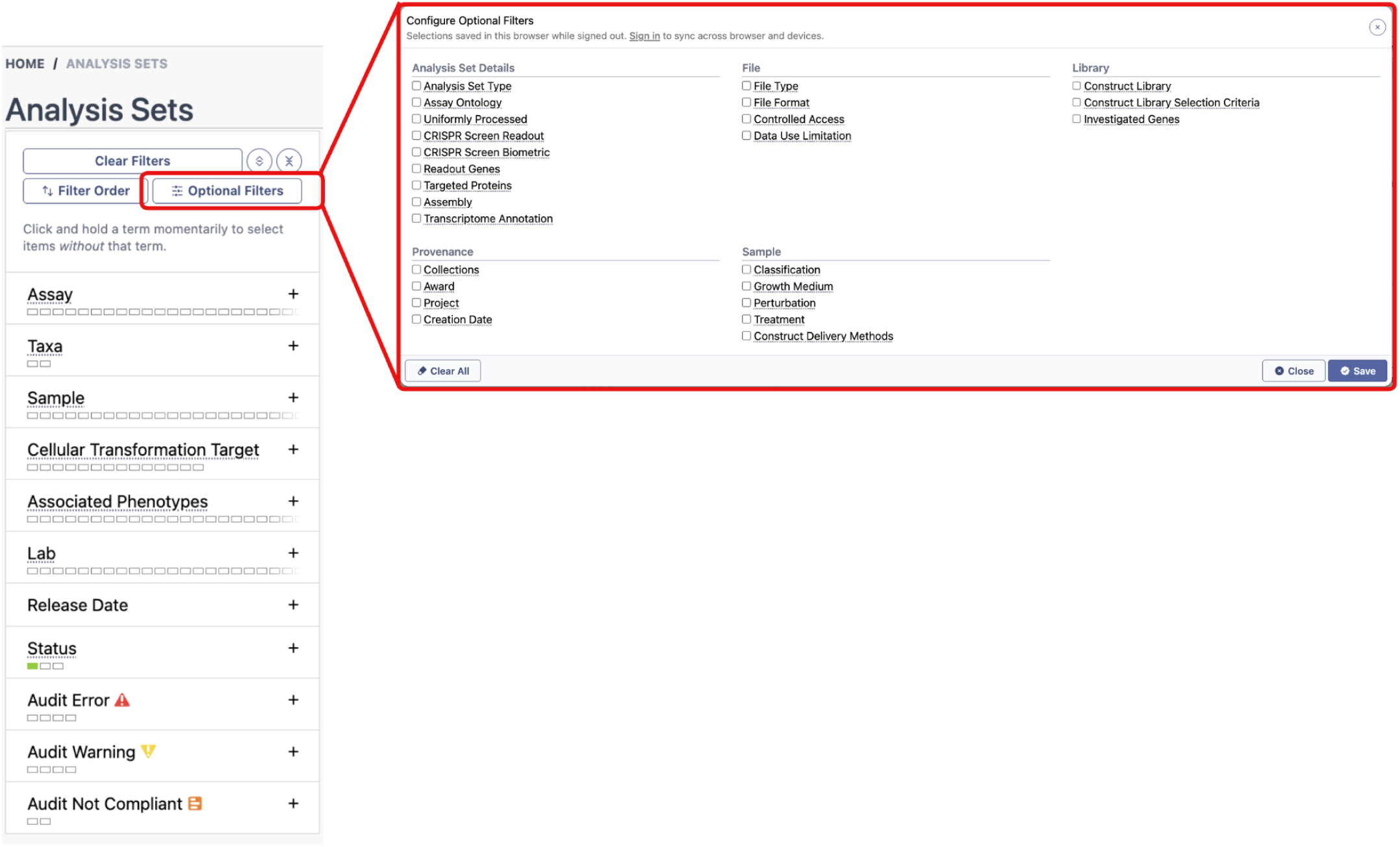
IGVF Data Portal facet overview. The IGVF Data Portal uses facets as a tool for users to filter for data of interest using the metadata associated. A screenshot of the sidebar on the portal search page for analysis sets, which has a list of “facets” used to filter search results according to different metadata fields. Each facet includes a hover over tooltip with an explanation of the facet. At the top of the sidebar are different facet options. The “Clear Filters” button will remove all currently selected filters from the search. The “Filter Order” button enters a special mode allowing users to drag and drop the facets in the sidebar to their preferred order. The “Optional Filters” button opens the pop-up menu displayed in red-lined inset, where users can select additional facets to display in the sidebar.

In addition to faceted search, the portal also supports a free text search box, which returns hits organized by object type. The search box functions similarly to a standard keyword search for quicker navigation when the precision of faceted searching is not required.

### Data Download

Downloading files is a key purpose of the IGVF Portal, which provides multiple download options to accommodate different user needs. Individual files can be downloaded directly from the file tables displayed on file set summary pages. Users can also download files in bulk using the download button at the top of a file search results page. Similarly, file set search results include a download button that allows users to download all files associated with the file sets returned by the search.

To download all files associated with a specific file set, the DACC has developed the batch download functionality. This portal application easily aggregates and downloads all related files, which are in most cases split up across multiple file sets. This feature automatically traverses upstream and downstream relationships between file sets to identify and aggregate associated files, providing users with a comprehensive download package. There are 3 options available for users to select which files should be included: (i) files in this file set, which includes all files directly associated with the selected file set; (ii) files in this file set and their input files, which additionally includes input files used to generate them; and (iii) files in this file set, their input files, and downstream analysis files, which further includes downstream analysis files derived from them.

As illustrated in **Supplementary Figure 1**, the diagram outlines 3 example scenarios of batch download depending on the selected file set and its position within the data model. The direction of file traversal (upstream, downstream, or both) depends on the selected download option. **(A) Upstream starting point —** The selected file set is located near the beginning of the workflow. Batch downloading can include its associated input files and downstream analysis files. **(B) Intermediate starting point —** The selected file set is located in the middle of the workflow. Batch downloading can traverse both upstream and downstream relationships to retrieve associated files. **(C) Downstream starting point —** The selected file set is located near the end of the workflow. Batch downloading can retrieve upstream input files and any downstream analysis files, while excluding unrelated branches of the workflow. These download options are available only for open-access files. Controlled-access files cannot be downloaded directly from the IGVF Portal and must instead be accessed from AnVIL dbGaP authorization.

### Integration of Portal and Catalog

All data loaded into the IGVF catalog comes from tabular files that are stored on the IGVF Data Portal for provenance. These data include novel results generated by the IGVF Consortium as well as pre-existing data resources, such as enhancer-gene regulatory interaction predictions from ENCODE **[6]**, eQTLs from EBI eQTL Catalogue **[7]** (https://www.ebi.ac.uk/eqtl/), protein complexes from EBI **[8]** (https://www.ebi.ac.uk/complexportal/home), variant annotations from FAVOR database **[9]** (http://favor.genohub.org/) and more, which provide additional information and context for IGVF results. Metadata fields indicate which files are included in the catalog, as well as the specific catalog version, so that the exact source files can be found on the portal. For example, the files containing data included in v1.3 of the catalog are available at https://data.igvf.org/search/?type=File&collections=IGVF_catalog_v1.3.

The DACC works with IGVF labs to review incoming data and understand how the data best fits into the knowledge graph. Scripts are then written to load the data sets (https://github.com/IGVF-DACC/igvf-catalog), including essential metadata such as biosamples and assay types curated from the IGVF Data Portal. By modeling all data points in one knowledge graph with their biological context, the IGVF Catalog provides users a resource to query by genetic variants, genes or genome coordinates, and retrieve integrated experimental evidence and computational predictions from various data sources. In total, the knowledge graph contains over 3 billion nodes and 8.6 billion edges, stored in an ArangoDB graph database (https://arango.ai). Users can query the data from the IGVF Catalog’s API (https://api.catalogkg.igvf.org/) as well as a web interface (https://catalog.igvf.org). Further details about the catalog are described in the upcoming Catalog publication (Hitz *et al*, in preparation.)

### Uniform Data Processing

In collaboration with IGVF focus groups and labs, three pipelines were developed to process IGVF data in a uniform manner: the single cell RNA-seq/ATAC-seq pipeline by the single cell focus group, the MPRA pipeline by the MPRA focus group, and the Perturb-seq pipeline by the CRISPR focus group. Of these three, the DACC oversees the operation of the single cell pipeline directly using Google Cloud and Terra.bio, while the MPRA and Perturb-seq pipelines are operated separately by IGVF labs. These pipelines are developed to integrate with the data portal by retrieving raw data from the portal, and then depositing the processed results back to the portal, with metadata to track the relationship of the files. Uniformly processed datasets and data files on the IGVF portals are marked with “Uniformly Processed” badges.

#### Single cell pre-processing pipeline

To facilitate integrative analysis across single-cell experiments, the IGVF Single Cell Focus Group developed a portable WDL pipeline, deployed on Terra.bio (https://terra.bio/), to uniformly process raw FASTQ files (https://github.com/IGVF/single-cell-pipeline, publication in preparation). The pipeline uses kb-python to quantify RNA-seq reads and Chromap to align ATAC-seq reads. QC metrics are also generated for each output. Supported assays include standard 10x Multiome, Parse SPLiT-seq, and SHARE-seq **[10,11]**. The DACC also developed a pipeline management tool (https://github.com/IGVF-DACC/sc-pipeline-management) that extracts the required input parameters from the data files and metadata submitted to the portal and launches the pipeline run. The same tool then accessions the output files back to the IGVF portal, with QC metrics stored as quality metric objects linked to each file. Once a lab’s data submission is complete, the DACC runs the pipeline and submits the outputs, a process that is largely automated. The output of the pipeline can be used in downstream analysis and pseudobulk annotation.

#### Sequence specification documents

The IGVF Data Portal requires a machine-readable specification, called seqspec, to document accurate sequencing library structures **[12]**. The seqspec records the FASTQ read files, content of read regions, and relevant metadata using a standardized and controlled schema. Single cell sequencing and Perturb-seq data rely on accurate seqspec YAML files to automatically and reproducibly generate input parameters for the uniform pipelines. Other non-single cell studies are required to submit a seqspec styled document detailing the libraries as references. The IGVF requirement for documentation of the library structure helps ensure the longevity and reusability of the sequencing data by future researchers.

#### MPRA pipelines

IGVF Consortium and the MPRA focus group developed the IGVF MPRA uniform pipeline (MPRAsnakeflow, https://github.com/kircherlab/MPRAsnakeflow), which creates count tables for candidate sequences using MPRA FASTQ and library design files **[13]**. The IGVF MPRA quantification pipeline (https://github.com/kircherlab/IGVF_MPRA_quantification) was developed to quantify MPRA reporter expression, incorporating the standardized input and output file formats established by the MPRA focus group **[14]**. The pipeline supports quantification of both variant- and element-level regulatory activity, facilitating consistent and reproducible analysis of MPRA datasets.

### Programmatic Access

Programmatic access to the IGVF Data Portal is available by making requests to https://api.data.igvf.org/ via a Hypertext-Transfer-Protocol-(HTTP)-based, Representational-state-transfer (RESTful) application programming interface (API). Responses are provided in standard JavaScript Object Notation (JSON) format. Many scripting languages, such as Python or R, have pre-existing libraries to handle JSON format, allowing user flexibility in how they choose to programmatically interact with the portal. The DACC has also made a Python client available at https://github.com/IGVF-DACC/igvf-python-client with methods to search and retrieve data from the portal. Documentation and examples on usage of the API is available at https://data.igvf.org/help/igvf-api-spec/.

### Documentation and AI Agent Support

To help users navigate the data portal, the DACC has created documentation describing the data model, UI features, and more, which can be found in the Help section of the portal. The General Help provides overviews of ontologies, common file formats and content types of files, status terminology, programmatic REST API usage, and more. The Data Submission section includes documentation targeted towards data submitters regarding submission tools and how to prepare data for submission.

In addition to documentation pages, the DACC has worked on incorporating hover-over tooltips throughout the portal UI, providing a quick and convenient method to see explanations for unfamiliar assays, file types, samples, and so on. Furthermore, multiple IGVF labs have started integrating these available tools to create AI agents for data submission, query, and visualization. The DACC has also set up an IGVF portal MCP to assist in these efforts, available at https://github.com/IGVF-DACC/igvf-portal-mcp.

## Future Directions

Although the IGVF Consortium has made much progress in exploring how genomic variation affects function, more remains to be studied. The DACC will continue updating the data model to accommodate data from new and cutting edge assays as well as novel computational methods. The DACC also seeks to expand collaborations with other consortia and other outreach efforts to publicize IGVF data. At the same time, the DACC will design new UI features to improve data discoverability, searchability, and accessibility, such as by highlighting key resources and featuring them on the home page, based on feedback from members of the scientific community. Furthermore, the DACC will optimize and scale the database backend to ensure stable access as the total amount of IGVF data increases and as programmatic, agentic data retrieval becomes increasingly common. The DACC will continue updating the IGVF portal to meet evolving needs, ensuring the longevity of the IGVF data resource.

## Supplementary Figures

**Supplementary Figure 1:**
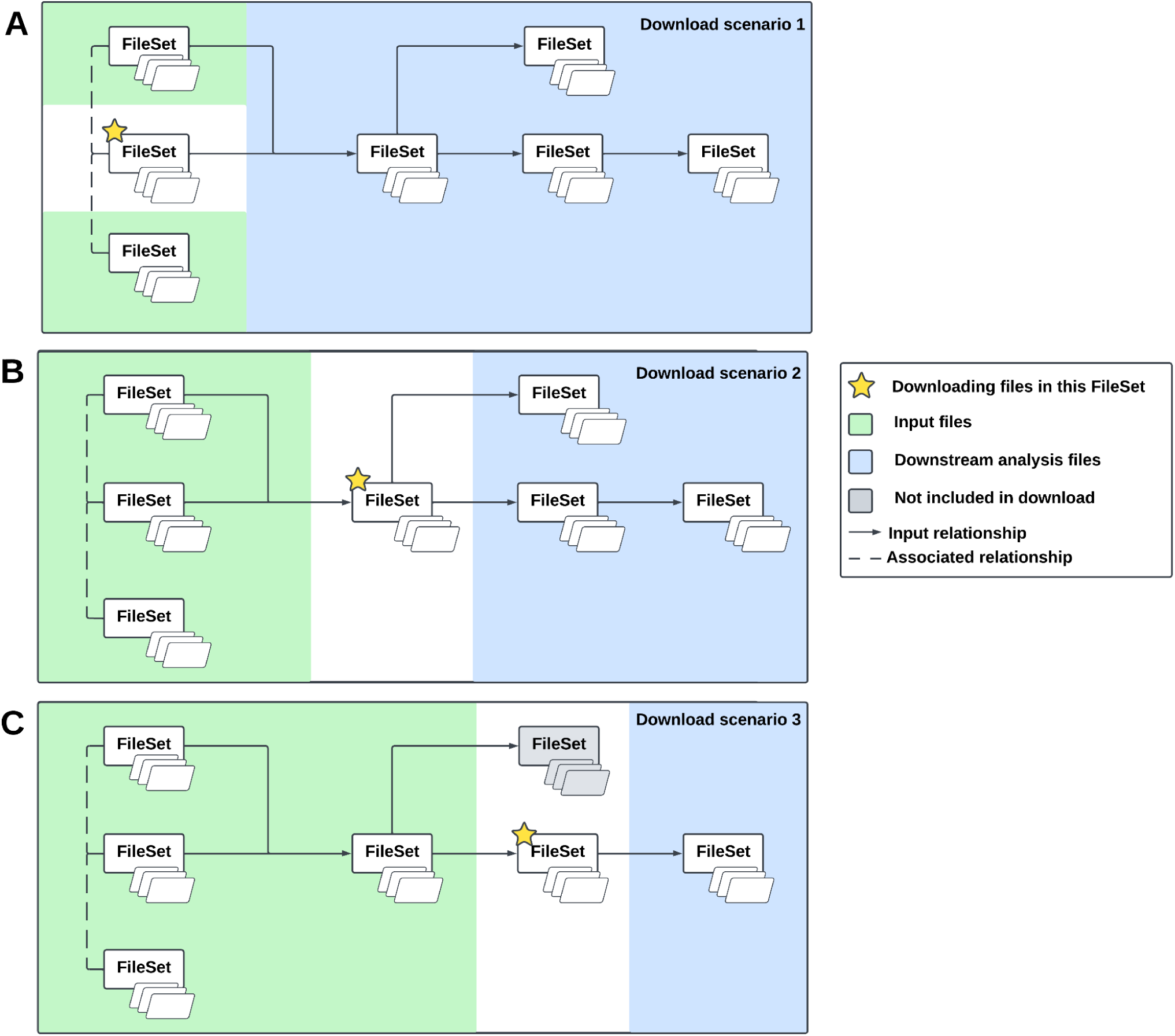
Overview of IGVF Portal data batch download. This diagram demonstrates how the IGVF Portal identifies and retrieves related files when batch downloading is initiated from a file set at three different positions in a data model: **(A)** upstream, **(B)** intermediate, and **(C)** downstream. The selected file set is indicated by a yellow star. Green regions represent upstream input files, blue regions represent downstream analysis files, and gray regions indicate files excluded from the download. Solid arrows indicate input relationships between file sets, while dashed lines indicate associated relationships. The batch download functionality traverses these relationships to identify relevant files, with the scope of included files varying depending on the selected starting point.

## Acknowledgements

This work was supported by the NIH NHGRI IGVF Program (U24HG012012). We thank members of Stanford IGVF DACC for developing and maintaining the IGVF Data Portal and IGVF Catalog as well as supporting IGVF Consortium member labs in data submission related functions. We thank all the IGVF Consortium members and institutions who have contributed to their experimental and predictive studies. Finally, we want to thank the IGVF Catalog team at Washington University in St. Louis for their contribution to the IGVF Catalog UI/UX development.

